# Pooled population resequencing of clam shrimp (*Eulimnadia texana*) from different vernal pools reveals signatures of local adaptation

**DOI:** 10.1101/223602

**Authors:** James G. Baldwin-Brown, Anthony D. Long

## Abstract

Vernal pool clam shrimp (*Eulimnadia texana*) are a promising model due to ease of culturing, short generation time, modest genome size, and obligate desiccated diapaused eggs. We collected Illumina data (Poolseq) from eleven pooled wild vernal pool clam shrimp populations. We hypothesized that restricted gene flow between vernal pools, separated by distances of 0.36 to 253 km, in concert with Poolseq data from each population, could be used to identify genes important in local adaptation. We adapted *Bayenv2* to genome-wide Poolseq data and detected thirteen genomic regions showing a strong excess of population subdivision relative to a genome-wide background. We identified a set of regions that appear to be significantly diverged in allele frequency, above what is expected based on the relationships amongst the populations. Regions identified as significant were on average 9.5 kb in size and harbored 3.8 genes. We attempted to identify correlations between allele frequencies at each genomic region and environmental variables that may influence local adaptation in the sequences populations, but found that there were too many confounding environmental variables to draw strong conclusions. One such genomic region harbored an ortholog of *Drosophila melanogaster* CG10413, a gene predicted to have sodium/potassium/chloride activity. Finally, we demonstrate that the identified regions could not have been found with less powerful statistics, i.e. *F_ST_*, or with a less contiguous genome assembly.

## Introduction

The clam shrimp *Eulimnadia texana* has, along with other vernal pool shrimp, been noted for its unique sex determining system (Sassaman and Weeks 1993), its rare (in Metazoa) requirement to reproduce via desiccated diapaused eggs (Sassaman and Weeks 1993), and its unique habitat. Further, *E. texana* is a relative rare androdioecious (Sassaman and Weeks 1993) species having three common arrangements of sex alleles (Sassaman and Weeks 1993) or “proto-sex chromosomes” (Weeks et al., 2010). The ability of eggs to remain in diapause for years at a time (Brendonck 1996) is especially valuable to geneticists because very few macroscopic animals exist for which populations can be archived for long periods without changes occurring in the genetics of the population (genetic drift, loss of linkage disequilibrium, etc.). In another paper we carried out a highly contiguous genome assembly of <u>*E. texana*</u> that resulted in a genome assembly with an N50 of 18Mb (Baldwin-Brown et al, in review), the availability of a high quality reference genome allows us to ask novel questions in this system. In this regards, an interesting aspect of *E. texana* is that they live in isolated vernal pools in the desert southwest of the U.S.A. The presumably naturally limited migration from pool to pool makes *E. texana* well-suited to the study of populations evolving in relative genetic isolation.

Various methods (Frichot et al. 2013; Günther and Coop 2013; Weir and Cockerham 1984; Nielsen et al. 2005; Voight et al. 2006) have been proposed for identifying signals of selection in wild populations. A higher *F_ST_* value than expected at some locus conditional on some sort of genome-wide average indicates that a force outside of genetic drift and migration is acting upon variation in an area of the genome (Akey et al. 2002). Recently, several analogous statistics, including *Bayenv2’s* Bayes factors (Günther and Coop 2013) and *LFMM’s* z-values (Frichot et al. 2013), have been developed that used Bayesian statistical methods to identify the likelihood or probability that a given polymorphism’s pattern of allele frequencies across sub-populations is explained by the genome-wide shared ancestry of the populations versus some sort of local adaptation. These statistics have an advantage over raw *F_ST_* in that they account for existing relationships between populations. Although there is no perfect method for detecting selection in wild populations, several studies (Lotterhos and Whitlock 2014) indicate that *Bayenv2* and *LFMM* are especially powerful for this type of analysis.

Pooled population sequencing (Poolseq) has been in use essentially since the advent of next-generation sequencing (Burke et al. 2010; Futschik and Schlötterer 2010). Poolseq allows for relatively inexpensive estimation of genome-wide allele frequency differences between populations. Here we use pooled population sequencing of shrimp from several pools to carry out population genetics analyses of this species, allowing us to identify common polymorphisms and compute classical population genetics parameters such as *Θ* and *ϱ*. We further used these POOLseq data to estimate *F_ST_* (Weir and Cockerham 1984), *Bayenv2*’s (Günther and Coop 2013) *X^T^X* and Bayes factors, and *LFMM*’s (Frichot et al. 2013) z-values to identify regions of the genome that have differentiated due to local adaptation. We determine *Bayenv2* to be the most appropriate method for detecting local adaptation from POOLseq data and a high quality reference genome, and identify 13 regions displaying a signature of local adaptation. We further identify correlations between these regions important in local adaptation and various environmental variables that differ between vernal pools. In particular, the collection date, male frequency, latitude, pH, inbreeding coefficient of the population, presence or absence of fairy and *Triops* shrimp, and elevation stand out as environmental variables that help explain population differences.

## Methods

### Shrimp collection and rearing

Clam shrimp populations were sampled from New Mexico and Arizona as previously described (S C Weeks and Zucker 1999). We acquired 11 soil samples, each from a different natural clam shrimp pool, to grow shrimp for sequencing (Figure 1, Supplementary data table 1); additionally, we sequenced one laboratory population (EE) that is directly descended from the WAL wild population, but has been reared in the lab for six generations. This population was derived from 265 WAL wild individuals, and was maintained at a minimum population size of 250 individuals for 6 generations. We hydrated the soil samples, and then collected 100 individuals (males and females) from each population on day 10 of their life cycle. These particular clam shrimp populations were chosen because ecological data were already available for these sites (Supplementary data table 1). Clam shrimp populations were reared in 50X30X8 cm disposable aluminum foil catering trays (Catering Essentials, full size steam table pan). In each pan, we mixed 500mL of soil with 6L of water purified via reverse osmosis. 0.3 grams of aquarium salt (API aquarium salt, Mars Fishcare North America, Inc.) were added to each tray to ensure that necessary nutrients were available to the shrimp. Trays were checked daily for non-clam shrimp, especially the carnivorous *Triops longicaudatus*, and all non-clam shrimp were immediately removed from trays. We identified the following non-clam shrimp: *Triops longicaudatus*, *Daphnia pulex*, and an unknown species of *Anostraca* fairy shrimp.

**Figure 1:**
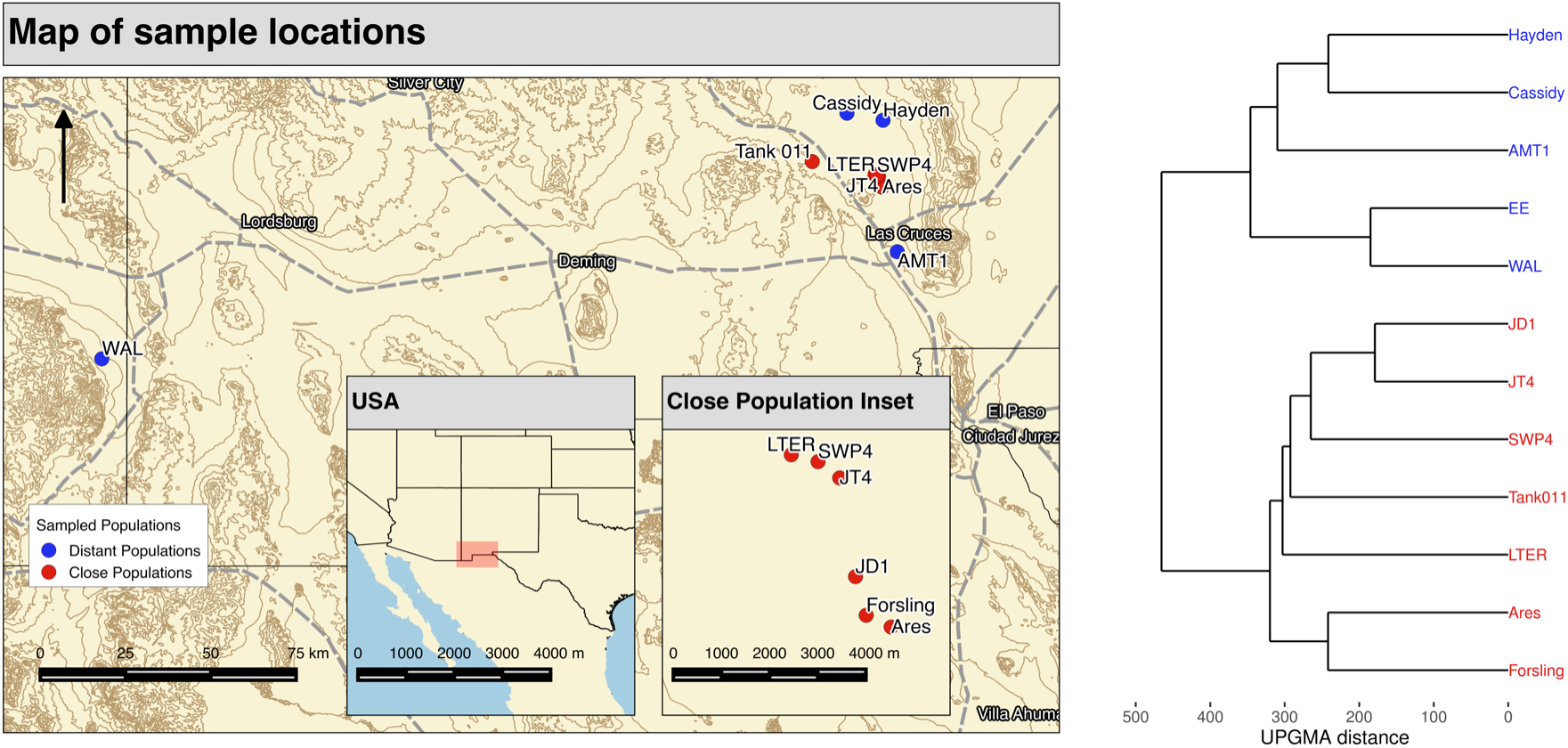
A map of the sampling locations for the 11 study populations and a UPGMA tree depicting the relatedness of the populations based on genome-wide allele frequency estimates. Colors correspond between the map and the tree. All populations were taken as soil samples from field sites in New Mexico and Arizona. The “EE” strain is a laboratory strain descended from WAL.

### Library preparation and sequencing

#### Illumina libraries

We produced our gDNA libraries using the Nextera Library Preparation Kit. We collected 100 random individuals from each population and pooled the individuals from each population to make each of the 12 libraries (one library per population). 13 cycles of PCR were used during the Nextera protocol, except in the case of the LTER and Tank 011 populations, where 15 cycles of PCR were used due to low yield. Each library was barcoded (Sup. table 1). Equal aliquots of each library were pooled, and the pooled samples were size selected on a Pippin (Sage Science, Beverly, MA) size selection instrument. The pooled libraries were sequenced over four runs of paired end 100bp Illumina sequencing, producing a total of 127Gb of data, or 844X of coverage. Full coverage statistics for each library are included in supplementary table 2.

### Comparison of wild populations

#### Genome assembly and annotation

We used the genome assembly and annotation generated by Baldwin-Brown et al. 2017 (in review). The genome assembly and annotation are available, along with all other scripts and major data, at the URL listed under “Data Availability”.

### Data preparation

Our pipeline for cleaning Illumina sequencing data, aligning to the reference, and calling SNPs was as follows: align using *BWA* (Li and Durbin 2009), deduplicate data using *Picard tools* (https://sourceforge.net/projects/picard/), and call SNPs using *GATK* (McKenna et al. 2010). After SNP calling, we censored SNPs by coverage using the following protocol: after merging the WAL and EE populations, remove all SNPs that have a mapped coverage of less than 10 or more than 200 in any population (in the two deeply sequenced samples, the 200 cutoff was applied to the coverages after random down-sampling of reads to match the less well covered populations), and remove all SNPs that, in any population, have a coverage more than 3 standard deviations from the population’s mean coverage. We performed this censoring separately for each of the three population comparisons examined in the results section. This removed a variable number of SNPs from the population depending on the coverages in each comparison, leaving a total of 1.4 million SNPs for further analysis in the full 11-population comparison. Command line options for *Picard tools, BWA*, and *GATK* are included in the supplementary texts, and the full scripts are available at GitHub (see Data Availability).

### Calculation of population genetics statistics

Our simple, genome wide estimate of *Θ* was calculated as:

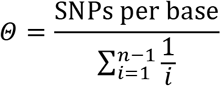

Where *Θ* is Watterson’s theta (Watterson 1975) and *n* is the sample size (approximately the average coverage of the genome). We calculated the genome-wide average *Θ* per basepair using the entire dataset independently for each of our populations, and then averaged the result to produce our reported *Θ* value. Note that Watterson’s estimator will be inherently biased in cases where SNP ascertainment is imperfect, an acute problem with Poolseq datasets. Rare alleles are likely to be underrepresented if SNP detection employs some sort of frequency cut-off in the pool (e.g. alleles at a frequency of <2% in the pool), but rare alleles will be overestimated if no cut-off is employed (and Illumina sequencing errors are erroneously called as SNPs). We accounted for this error by fitting the neutral site frequency spectrum expected by chance (Fu 1995) to all SNPs with a minor allele frequency above 0.1 (given our observed coverages, sequencing errors will only very rarely reach a frequency of greater than 10%), and then using that projected allele frequency spectrum to identify the expected number of true SNPs in our sample. Fu 1995 notes that the expected minor allele count, for coverage *n* and minor allele count class *i*, is equal to *ϕ*_i_*Θ*, where *Θ* is an estimator of 4*N_e_μ*, and

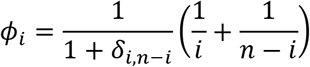

Here, *δ_i,n-I_* represents Kroneker’s delta, which is equal to one if *i*=*n*-*i*, and is otherwise equal to zero. For each population, we computed the expected *ϕ*_i_ based on the average coverage of that population, and then found the value of *Θ* that minimized the difference between the observed and expected number of SNPs with a minor allele frequency greater than 10%. From there, we multiplied the observed number of SNPs with a minor allele frequencing greater than 10% by 1/[the expected proportion of SNPs with a minor allele frequency of greater than 10%] to arrive at a project SNP count.

We calculated *ϱ* by first estimating the short-distance linkage disequilibrium using *LDx* (Feder, Petrov, and Bergland 2012). We then estimated *ϱ* by modeling decay of linkage disequilibrium (*r*^2^) with distance in basepairs using a non-linear model, as in (Marroni et al. 2011) (supplementary text). See results for more detail on *ϱ*.

### Fourfold Degenerate Sites

We generated a custom Python script for identifying fourfold degenerate sites for use in *Bayenv2*. This script identified sites based on the codon contents of the CDS in all *Augustus*-identified candidate genes. Candidate sites are not considered fourfold degenerate if even one transcript disagreed with that assessment. Fourfold degenerate sites were used for *Bayenv2*’s covariance matrix generation step because fourfold degenerate sites have been shown to be under selection less often than any other class of genomic site (Z. Yang and Bielawski 2000).

### Identifying differentiation

We calculated *F_ST_* via the Weir and Cockerham method (Weir and Cockerham 1984) using a custom *R* script. We calculated pairwise *F_ST_* for each pair of wild populations, and then reported the mean at each locus. We also calculated Bayes factors using *Bayenv2* (Günther and Coop 2013) both for population differentiation and ecological factor correlation. We did not use *Bayenv2*’s option to incorporate pooled sequencing variation into the Bayes factors because we could not get the program to finish when using that setting (supplementary text). In the case of population differentiation, we did not use the pooled sequence option because it hasn’t been correctly formulated (Günther and Coop 2013).

### Fitting Q-Q plot distributions

We fitted gamma distributions to the genome-wide collection of each of the 11-populations differentiation statistics (*F_ST_*, the *Bayenv2* and *LFMM* statistics, and *SweeD*’s CLR values) using *R*’s *optim* function. We allowed up to 1000 iterations of fitting and ensured that this limit was above what was needed for convergence in all cases. The allowed range for each of the fit parameters was [0.1,300]. We censored the bottom 10% and top 20% of data points from each distribution for the purposes of fitting in order to test whether observed extreme values would match the distribution obtained purely from the center of the distribution. *optim* minimizes a computed summary statistic for the fit of the distribution; in our case, we minimized the sum of the squared deviations between the observed and expected cumulative density statistics. We did not fit the lower tail of the observed data because some of the statistics have a mathematical lower bound not easy to accommodate with the gamma distribution (e.g., *F_ST_* can take on negative observed values); more importantly, we also did not fit the upper tail of the observed data, as this is where we expect loci showing an excess of differentiation to diverge from the null distribution describing the remainder of the data.

### Sliding window calculations

We performed windowed analyses by averaging values across SNPs with all statistics except *LFMM*’s *z*-values, which were combined in windows using the Fisher-Stouffer method, as detailed in the manual for *LFMM* (Frichot et al. 2013). We note that the Fisher-Stouffer method assumes independence of the values of the combined statistics, which is not necessarily true in this case due to linkage disequilibrium, so the *p*-values produced by this method should not be regarded as exact. In any case, we largely disregard *LFMM*’s results due to coverage as discussed below, and window averaged statistics are not used in any of the values reported here, but are merely used in plotting where indicated. We used 25-SNP windows in all cases except *LFMM*, where 100-SNP windows were used. We chose all of these based on visual examination of the statistics – these window sizes appeared to reduce noise while making peaks more visible.

### Significance

We consistently use a genome-wide false positive threshold of 0.05 where possible, (i.e., with *LFMM* and *SweeD*). With *X^T^X* and Bayes’ factors, computational limitations led to a significance threshold based on 4 million SNPs, or a significance threshold of 0.36; thus, the significance thresholds should be taken with a grain of salt. We generated this neutral distribution using simulation machinery from Gautier 2015 (Gautier 2015). This machinery uses a covariance matrix produced by *Bayenv2*, plus information on sampling and coverage, to generate the distribution of allele frequencies that we would expect if the sequenced populations are evolving neutrally and are related in the way that the covariance matrix describes. We generated 20 times the number of SNPs assayed throughout this paper (about 20 million) and used the maximum value as our significance threshold, giving a false detection rate of approximately 1 in 20, or 0.05. In the case of *X^T^X* and Bayes factors, we used the maximum value of 4 million SNPs.

In the case of 25-SNP windows we cannot use the simulation machinery of Gautier to generate a null distribution. This machinery simulates independent SNPs conditional on the among population covariance matrix, but sliding windows in real datasets are typically over SNPs in LD with one another. As a result, the 25-SNP average for real data has a much larger variance than the same statistic calculated over independent SNPs; hence, much of the genome would be interpreted as “significant” by a sliding window calculation. To obtain a significance cut-off for 25-SNP window averaged *X^T^X*, we identified by eye the location in a Q-Q plot (see above for Q-Q plotting details) where the empirical values diverged from the line of best fit, and used that as a significance threshold. This “inflection point” from the Q-Q plot method generally agrees with the single SNP threshold obtained via simulation.

### *BLAST* annotation

We annotated all gene functions using *blastp* to align the *E. texana* genes to the *D. melanogaster* NCBI protein database, and vice versa. We regard the mutual best hits (those pairs that had e-values below 10^-5^ in both directions, and that paired in both *BLAST* directions) as the annotations in which we were most confident. In the 13 peaks of high interest discussed below, we annotated the genes that did not have mutual best BLAST hits in *Drosophila melanogaster* by taking the most significant BLAST hit for each gene (identified using *blastp* against the *D. melanogaster nr* protein database) and assigning that putative identity to the gene of interest.

### Environmental variable descriptions

Some measured environmental variables require special description. “Date” is the date of collection of the soil. “Percent males” refers to the fraction of individuals that were male in hydrated samples. Surface area and volume were calculated based on measurements taken on-site at these pools. *Streptocephalus mackeni* and *Thamnocephalus platyurus* refer to the presence or absence of these species of *Anostraca* fairy shrimp, and “Fairy shrimp” refers to all fairy shrimp where the species was unknown

## Results

### Population Genetic Statistics

We collected pooled population sequencing data from our 12 populations (11 natural populations, and 1 lab population, EE, descended from the WAL natural population) (Fig. 1), calculated allele frequencies at each SNP using *GATK* (McKenna et al. 2010), and used the resulting allele frequency estimates for subsequent analyses. We first used a simple hierarchical clustering tree to look at genome-wide relationships among the populations (Fig. 1). The populations EE and WAL, being directly related by only 6 generations of laboratory maintenance, should be quite similar to one another; many of the natural populations appear to be as closely related to each other as EE and WAL are and even the most distant populations are less than twice as diverged. This observation contradicts the conventional wisdom that vernal pool clam shrimp populations rarely exchange migrants.

Supplementary Figure 1 plots the observed minor allele frequency spectrum and the expected allele frequency spectrum under neutrality for a *Θ* that matches the frequency distribution for SNPs at a frequency greater than 10% (see the methods for a description of this correction). Although we calculated *Θ* independently for each of our 12 sequenced populations, supplementary figure 1 displays the result produced if all alleles from all populations are aggregated for ease of viewing. The results in each population are qualitatively the same. The figure shows that the expected neutral allele frequency spectrum contains a large number of rare alleles that are failing to be identified by our SNP calling pipeline, almost undoubtedly because our pipeline includes an allele frequency cut-off to be considered a true positive SNP. We accounted for this underidentification of SNPs by re-computing *Θ* using only high frequency SNPs and inferring the existence of additional rare SNPs under the assumption of neutrality and Wright-Fisher sampling (Fu 1995). This produced an averaged over populations estimate of *Θ* per basepair of 0.00387. This value of *Θ* is fairly typical for invertebrates, and is close to the estimated *Θ* for *Drosophila melanogaster* of 0.0053 (Andolfatto and Przeworski 2001).

We used the result of *LDx* to estimate average linkage disequilibrium at various distances up to about 400bp, then identified the recombination rate based on decay of linkage disequilibrium according to (Marroni et al. 2011). We estimated the population average “rho”, *ϱ*, per basepair (the population-adjusted recombination rate per basepair) to be 0.0036 or a genomewide estimate of *ϱ* of 436,000. Under population genetic theory, the expected value of theta is equal to 4*N_e_μ*, assuming a mutation rate (*μ*) of 2.8×10^-9^ (the *Drosophila melanogaster* mutation rate per site per generation, from Keightley et al. 2014), we estimate the effective population size of the clam shrimp to be *Θ*/(4×*μ*) = 3.45×10^5^. Since the expectation of *ϱ* is similarly *4Nr* (where r is the size of the genome in Morgans), we estimate *r* to be 0.33 (or 33 cM). Assuming this estimate of *r* is correct, it seems quite low. An apparent map size of 33cM is consistent with a lack of recombination in amphigenic hermaphrodites, which can represent 80-90% of individuals in many clam shrimp populations (Weeks et al. 1999b), suggesting a male map size closer to 300cM, consistent with Drosophila. That said, our estimates of *ϱ* need to be interpreted with some caution, linkage disequilibrium was estimated using only short reads, and is thus only estimated out to ~450bp. It is conceivable that longer range LD paints a different picture and/or does not follow the predicted decay rate (W. G. Hill and Weir 1988).

### Genome-wide Selection Detection

#### Pairwise population differentiation comparisons

We initially compared the WAL and EE populations. Since the EE population is a direct descendant (6 generations in the laboratory at ≥250 individuals per generation) of the WAL population, there are several reasons that a pairwise comparison of WAL and EE is of interest. First, if minimal differences between WAL and EE exist, then they can be combined to increase coverage of WAL in the 11-population analysis. Second, if we observe substantial regional differentiation between these two populations, this would be strong evidence of selection due to domestication in the EE population (that is, selection due to being reared in laboratory conditions). And third, the level of differentiation between the WAL and EE populations, which have a known history, could inform inferences about the history and relatedness of the wild populations. Under neutrality, *F_ST_* is expected to be exponentially distributed with a lambda that can be calculated from the empirical *F_ST_* distribution (Elhaik 2012). We computed *F_ST_* for this pair of populations and used a Q-Q plot to compare the observed distribution of *F_ST_* to an exponential distribution; however, rather than generating the theoretical distribution by computing lambda based on mean *F_ST_* as in Elhaik 2012, we fit our exponential distribution the middle 70% of the data using *optim* in R. (Sup. Fig. 2, see methods). Q-Q plots have great utility for genome-wide datasets where one expects a summary statistic from the vast majority of the genome to have a distribution consistent a null hypothesis, with the statistics from a small subset of SNPs distributed differently. The vast majority of data points will fall along a straight line consistent with the quantiles of the null distribution, with larger values of the observed summary statistic deviating sharply upward from that same straight line (see McCarthy et al. 2008 and Pearson and Manolio 2008 for reviews of how to interpret Q-Q plots in the context of genome-wide association studies where they have had a large impact). In the case of *F_ST_*, the large majority of the SNPs should have *F_ST_* statistics distributed according to an isolation by distance model, with a small subset of SNPs exhibiting higher *F_ST_* statistics suggestive of local adaptation. Q-Q plots of *F_ST_* (Sup. Fig. 2, first panel) indicate that, while there is a small bias toward higher allele frequency differences between WAL and EE compared to the theoretical expectation, the deviation is small; however, the fit of an exponential in this case does not seem to be a very accurate representation of the shape of the data. Normal and gamma distributions, fit with and without logging, had equally poor fits (data not shown). We suspect part of the difficulty in fitting *F_ST_* statistics to a simple exponential distribution is due to observed *F_ST_* statistics occasionally being negative.

The second panel of supplementary figure 2 represents the same Q-Q plot arrangement as the first panel, but uses a 25-SNP-windowed average of *F_ST_* rather than raw *F_ST_*. In this case we fit a normal distribution in place of an exponential distribution, since the mean of many independent identically distributed exponential random variables converges on a normal (Stuart and Ord. Kendall’s Advanced Theory of Statisitics: Vol 1 Distribution Theory, 6^th^ ed. 1994. New York: Halsted Press). As with the first panel, the genome can be construed as falling on a mostly straight line with the exception of two points. Both of these points appear to be SNPs that have escaped quality filtering but are misleading: one resides on the edge of a small contig and is more than 200kb from the next non-censored SNP, while the the other appears to be one of only two non-censored SNPs on its 36kb contig. Since there is little evidence that the WAL and EE populations are different from one another, we chose to pool them for all subsequent analyses. It is noteworthy that the null distributions we tested do not seem to be excellent fits to the observed, and indeed we tried many other distributions and found that the fit was no better; we propose that this is due to the complex properties of the *F_ST_* statistic, which has been noted elsewhere to have maximum and minimum values that vary from locus to locus (Jakobsson et al. 2013).

We next examined the two high coverage lines, LTER and Tank011. These two populations were sequenced much more deeply than the others, and thus have much more accurate estimates of allele frequency at every SNP in the genome (i.e., 206X average coverage for LTER and Tank011 compared to 48X average coverage for the other nine populations). We generated Q-Q plots of *F_ST_* for this pair of lines according to the same scheme used to compare WAL and EE (Sup. Fig. 3). We had many of the same problems fitting single SNP *F_ST_* and sliding window *F_ST_* as with WAL and EE, with the fits exhibiting similar properties. Based on single SNPs (first panel) *F_ST_* statistics fall along a straight line, with the exception of values of *F_ST_* greater than 0.5 where the statistic seems to max out, suggesting very little signal at single SNPs. In comparison with the relatively subtle divergence from the expectation line observed with the WAL vs. EE *F_ST_* Q-Q plot, this plot shows a dramatic divergence at the upper end of the distribution, though it appears to level off as *F_ST_* approaches 0.5. Likewise, there is a much more dramatic upward trend in the 25-SNP windowed Q-Q plot for LTER vs. Tank 011 than for WAL vs. EE. Like the single SNP *F_ST_* distribution at very large values of average *F_ST_* the curve for the 25-SNP window also flattens, again likely due to the *F_ST_* statistic saturating. Overall there is some indication of greater than expected population differentiation for some regions of the genome. This being said, the power of *F_ST_* to detect differences between populations seems to be weak if both windowing and 200X sequence coverage is necessary in order to identify a signature of local adaptation (that *F_ST_* also saturates is an addition problem). We later show that *F_ST_* seems to do a poor job of identifying local adaptation when we consider all 11 populations, and is perhaps not the most suitable statistic for detecting local adaptation.

#### Site frequency spectrum derived inference of selection

From here, we moved on to analyzing all 11 populations together. The WAL and EE populations were pooled, and LTER and Tank011 were each down-sampled to ~48X coverage. We began by performing a scan for selection based on site frequency spectra using *SweeD. SweeD* identifies variation in the site frequency spectrum (specifically, a lack of rare alleles) and takes this as evidence of a selective sweep in the recent past. We ran *SweeD* both for each individual population and for all populations together, but owing to the similarity between the results, and the fact that we believe the results are not highly informative, here we present only the full 11-population result. We plotted *SweeD*’s composite likelihood ratio (CLR) using a Manhattan plot (Sup. Fig. 4). We know that the site frequency spectrum is not accurately represented by the SNPs that we have identified using Poolseq (see the section on population genetics and supplementary figure 1 for more detail), thus *SweeD* should produce many false positive calls under these conditions. Since rare SNPs are difficult to distinguish from errors, many rare SNPs are never called, leading to an allele frequency spectrum that is skewed toward common alleles. Because *SweeD* identifies selection by finding regions of the genome whose site frequency spectra deviate dramatically from neutrality, we should expect and do indeed find that a large portion of the genome appears be under selection. Thus, the results of *SweeD* should be treated with skepticism in this case, and perhaps in other cases where pooled sequencing data has been used to ascertain SNPs.

### Population differentiation in all 11 wild populations

We next carried out an analysis to identify population differentiation using all 11 natural populations. We chose to use *F_ST_* and *Bayenv2’s X^T^X* statistic to scan for differentiation. For Q-Q plotting, we generated a theoretical distribution by fitting a two parameter gamma distribution to the middle 70% of the empirical data, as described in the methods. Q-Q plots of experimental *F_ST_* vs. the exponential distribution (Fig. 2) do not demonstrate any evidence of population differentiation. The Q-Q plot for *F_ST_* suggests little signal that could be used to detect regions of local adaptation. On the other hand, *X^T^X* seems to identify a subset of SNPs (with log_10_(*X^T^X*) statistics greater than ~1.45; Fig. 2) as being involved in local adaptation. The increased power in *X^T^X* compared to *F_ST_* is perhaps unsurprising as it takes into account the level of relatedness amongst the populations. A simple UPGMA tree comparing the populations based on genome-wide allele frequencies (Fig. 1) indicates that several of the populations are very similar to each other. In fact, knowing that “EE” is directly descended from “WAL” and separated by only 6 generations of laboratory breeding, the UPGMA tree makes it evident that some of the populations are nearly identical in terms of allele frequencies. Thus, *X^T^X*, which takes into account the relationships between the populations, should perform better when attempting to identify differentiated loci. Additionally, when we compute a 25-SNP-window average of *X^T^X* (Fig. 2), we arrive at a qualitatively very similar result: strong linearity with the gamma distribution until log_10_(*X^T^X* is greater than approximately 1.40.

**Figure 2:**
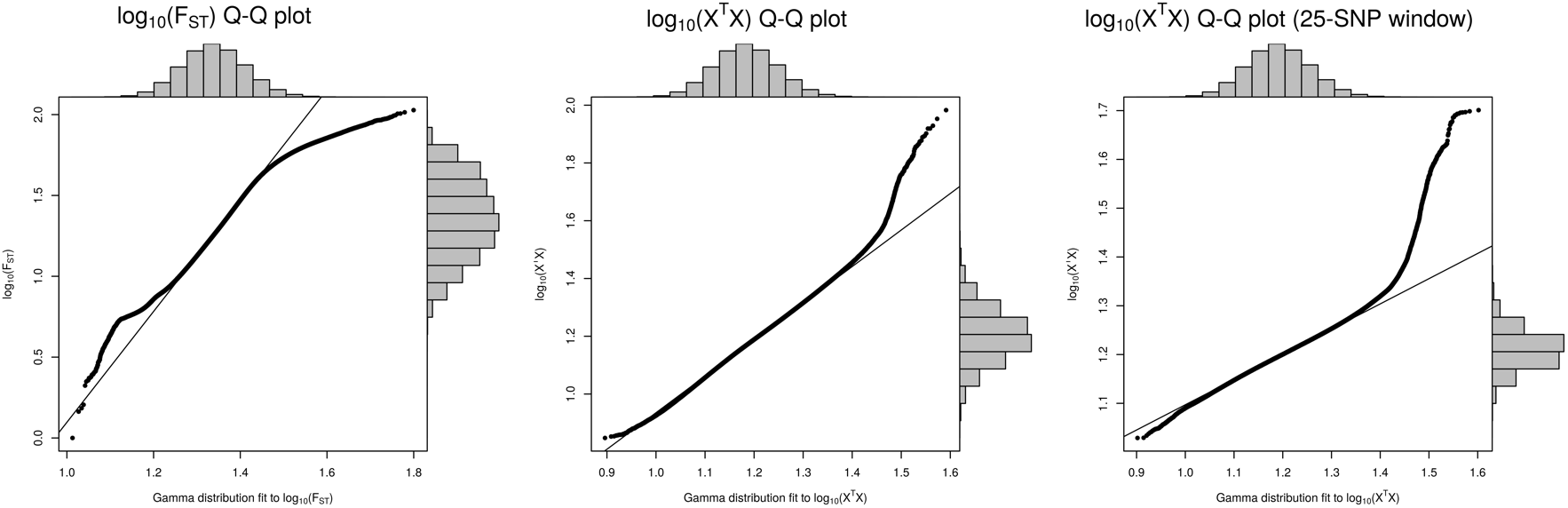
Quantile-quantile plots of two statistics used to identify regions with high population differentiation in the clam shrimp genome. Left: mean pairwise *F_ST_*, logged. Center: Bayenv2 *X^T^X*, logged. Right: Bayenv2 *X^T^X* 25-SNP-window average, logged. All three datasets are plotted against a two-parameter gamma distribution fit to the middle 70% of the observed data (methods). The straight line is a linear regression fit between observed and expected statistics for the same middle 70% of the data. The bar charts along the sides of the graph are histograms of the data along their respective axes. From the histograms of the observed statistics it is apparent that large departures from the line of best fit in the tails is driven by a very small fraction of the total loci.

We next generated Manhattan plots for single SNP *F_ST_*, single SNP *X^T^X*, and 25-SNP windowed *X^T^X* (figure 3). For *F_ST_* we set a genome-wide significant threshold from an exponential distribution with *L* = 1/mean(F_ST_). For single SNP *X^T^X* we used a simulated set of read count value as controls to set a genome-wide significance threshold of 0.36 (i.e., a single false positive SNP roughly every 3 genome scans), a more accurate determination of this threshold was largely dictated by computation limitations. The threshold of 37.7 (or log10(*X^T^X*) = 1.6) determined in this manner agrees with one obtained by simply taking the inflection point from the Q-Q plot of figure 2. For the 25-SNP windowed *X^T^X* analysis, SNPs in a window are not independent of one another, and we cannot simply take the average of 25 unlinked SNPs from our simulated read count experiment to set a threshold, as that would be anti-conservative. Instead we determined significance from the inflection point of the Q-Q plot at a log10(*X^T^X*) of 1.4. *F_ST_* (panel A) stands out here as having no peaks that indicate local adaptation.

**Figure 3:**
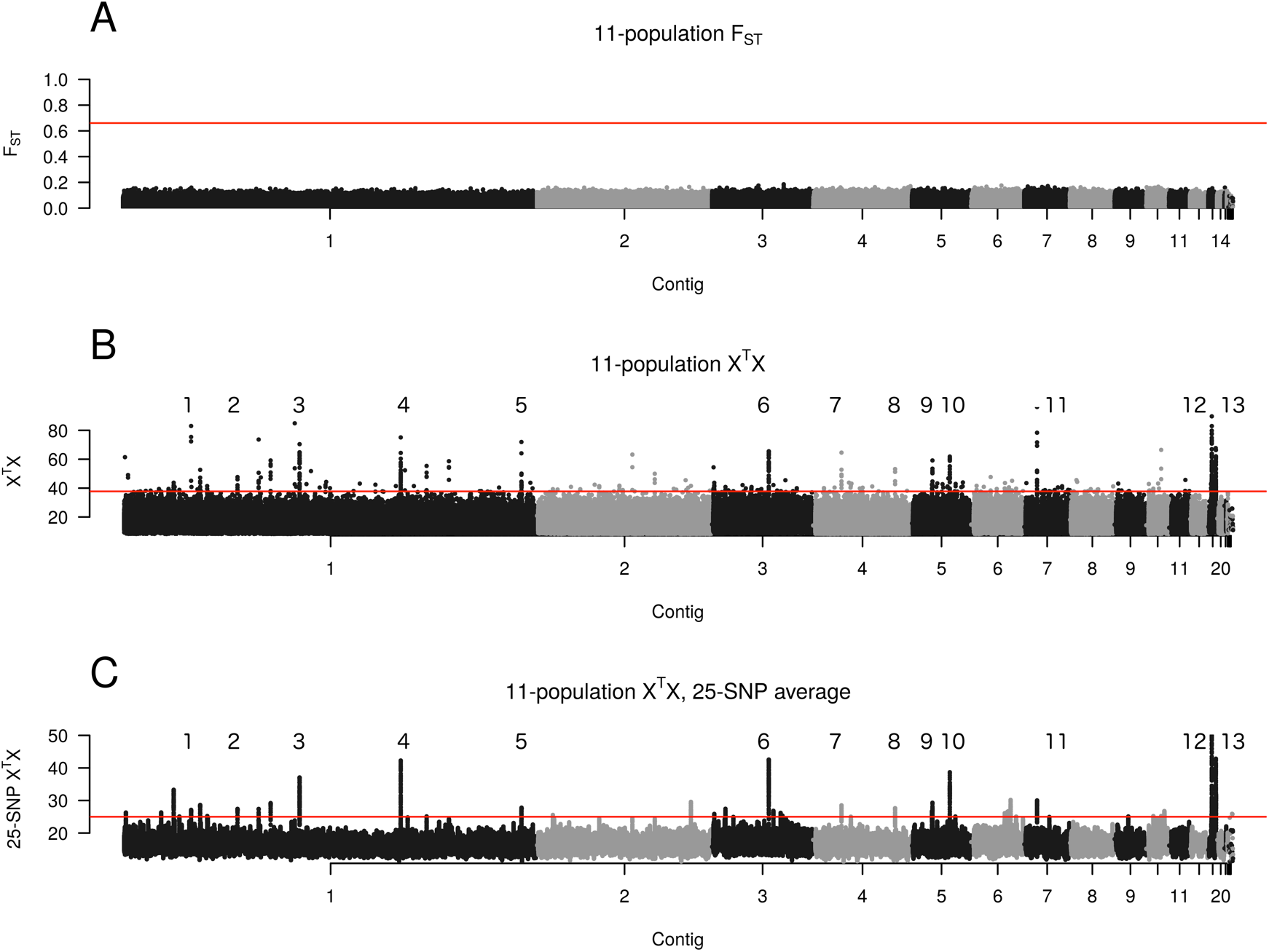
Manhattan plots of support for an excess of differentiation among the 11 populations. Plots are as follows: A: mean pairwise *F_ST_*; B: Single SNP Bayenv2 *X^T^X* statistic; C: 25-SNP window Bayenv2 *X^T^X statistic*. Identified peaks are numbered 1-13 and are referred to as such throughout this work. Red lines correspond to significance thresholds set as described in the methods.

On the other hand, the *X^T^X* statistic reveals numerous large peaks (plus a handful of smaller peaks) in the Manhattan plot of panel B. Upon closer inspection of these peaks, we noted that they were characterized by their contents: a subset of SNPs, scattered throughout a small region, with extremely high *X^T^X* values, and a relatively large number of SNPs with average *X^T^X* values (see Sup. Figs. 5-10). Based on this, we concluded that we could detect the regions with a large number of supporting polymorphisms by taking a power-transformed average of the data. Such an average would put extra weight onto high values, and could identify regions that contained large numbers of elevated values. We arbitrarily chose an 8^th^-power-transformed average of a 1000-snp window, with a cutoff (after back-transforming) of 24 (Sup. Fig. 13). We manually excluded regions at the edges of chromosomes, as well as regions that were embedded in overall high-*X^T^X* portions of contigs (i.e., those without a discernable peak), and those that were significant, but appeared to be so in only a few windows. See Sup. Fig. 13 and its description for details of the exclusions. From this, we identified 13 peaks that we number 1-13 in the Manhattan plot of panel B. These peaks extend well above our significance threshold, and consist of tens to hundreds of SNPs in regions small enough to contain one to three gene candidates. The 25-SNP sliding window average of *X^T^X* (panel C) seems to produce a result broadly consistent with the single SNP Manhattan plot, with some peaks becoming more pronounced (i.e., peak 4). We note that there is nothing special about the 25-SNP window size, as it was chosen arbitrarily based on the ability to resolve the peaks clearly, and qualitatively similar results are obtained with different windows. Since the windowed analysis does not seem to identify more peaks than the single SNP analysis we focus further discussion on the un-windowed *X^T^X* statistics.

We identify a total of 501 single significant SNPs, with 676 genes and 198 annotated genes (i.e., genes with a mutual *BLAST* hits to *D. melanogaster)* within 5kb. *GOrilla* (Eden et al. 2009) indicates that genes related to external visual stimuli are overrepresented in this set (Sup. Table 3). Of the 501 single SNPs above the significance threshold, 407 (or 81%) are associated with the 13 numbered regions, which appear as sharp peaks in the Manhattan plot. Further, the SNPs in the 13 numbered regions are associated with only 50 genes (22 being well annotated). Thus, based on the *X^T^X* test statistic, a small, localized fraction of the genome seems to account for much of the signal of local adaption. Figure 4 examines two such peaks in detail, and Supplementary Figures 5-10 the other eleven. Table 1 further details the boundary of each peak. From Figure 4, it is clear that resolution of these peaks is quite narrow, with peak widths in the range of 1-15kb, and most peaks containing less than 5 genes. We identified the probable identities of genes in the thirteen regions and compiled a table (Sup. Table 4) of these genes along with their exact locations relative to the peaks. Several peak associated genes have well-documented functions, including *Rumpelstiltskin*, *okra*, *Cp1*, *SNS*, *Dscam2*, *pyridoxal kinase*, and *Ublcp1*.

**Figure 4:**
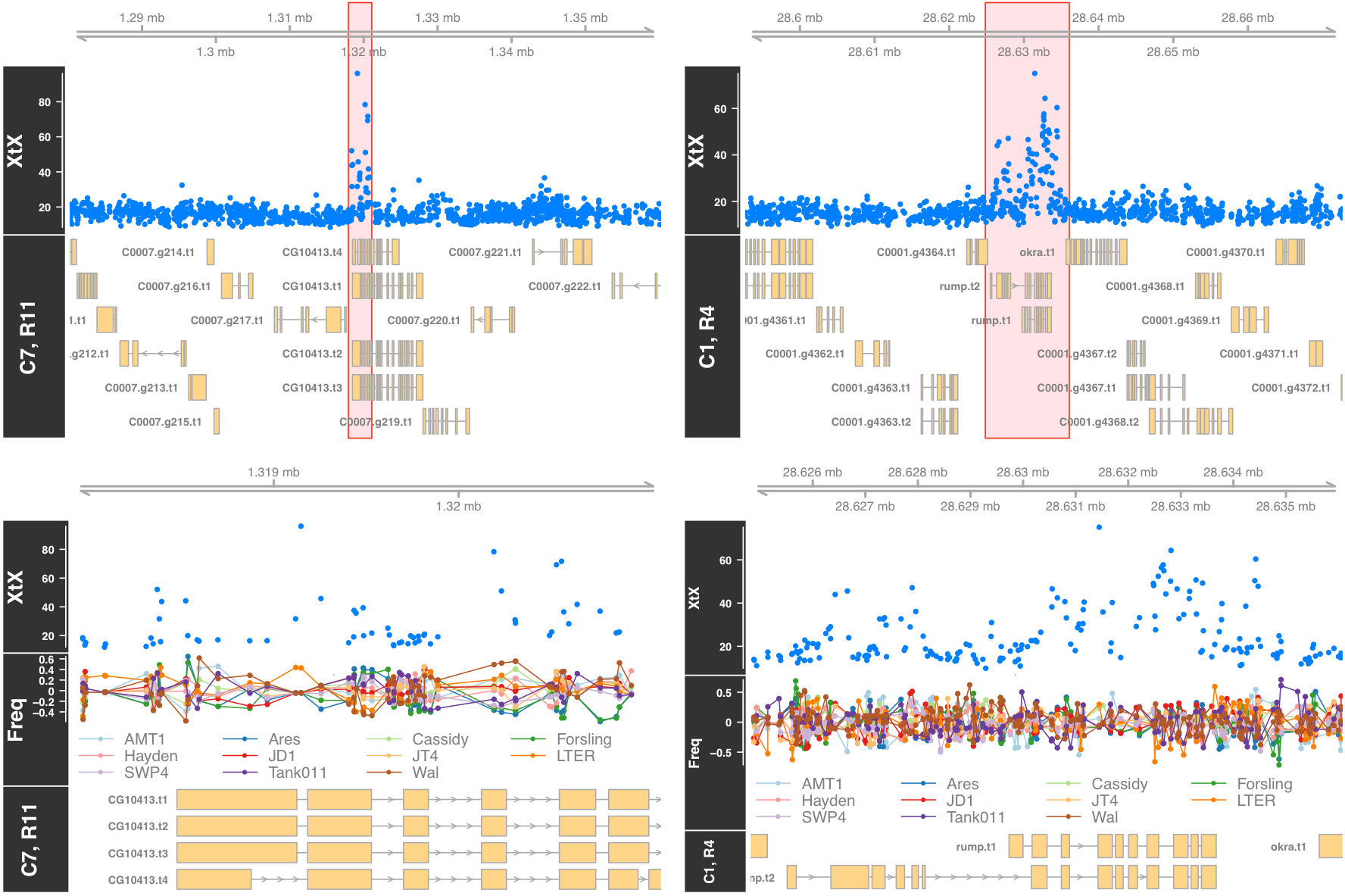
Manhattan plots of single SNP *X^T^X* values indicating excess differentiation among the 11 populations for the regions 11 and 4 as indicated in Figure 3. The plots indicate the signal is highly localized, often suggesting a single gene (CG10413 for locus 11, and rump or okra for locus 4). The red rectangle in each upper plot indicates the region shown in the corresponding lower plot. The “C” and “R” indicators in the titles indicate the contig and region number. “Freq” refers to the allele frequency in each population as a deviation from the mean for that locus.

**Table 1:** Major significant regions according to the 11 -way XT X population differentiation analysis

| Region | Contig | Range (bp) | Width (bp) | Maximum $X^T X$ |
| --- | --- | --- | --- | --- |
| 1 | 1 | 7960415:7961229 | 814 | 52.6524 |
| 2 | 1 | 11807825:11808199 | 374 | 47.6688 |
| 3 | 1 | 18205259:18209499 | 4240 | 70.4348 |
| 4 | 1 | 28626427:28634469 | 8042 | 75.102 |
| 5 | 1 | 41068868:41081593 | 12725 | 71.9472 |
| 6 | 3 | 5849714:5855825 | 6111 | 65.5048 |
| 7 | 4 | 2920923:2921932 | 1009 | 64.5828 |
| 8 | 4 | 8446194:8447185 | 991 | 53.1936 |
| 9 | 5 | 2113388:2117449 | 4061 | 59.1548 |
| 10 | 5 | 3903600:3908963 | 5363 | 61.8584 |
| 11 | 7 | 1318370:1320640 | 2270 | 96.1406 |
| 12 | 14 | 369627:420280 | 50653 | 89.70358 |
| 13 | 14 | 791617:817899 | 26282 | 67.7304 |

The apparent high localization ability of the *X^T^X* statistic in some cases suggests individual SNPs of importance. Within the 13 regions exhibiting peaks of significance several (*cf*. regions 1,2,5,7,8,9,11) are narrow enough that one or a small number SNPs stand out as being much more significant than their neighbors (Fig. 4, Sup. Figs. 5-10). We noted a single, highly significant SNP in regions 1, 2, 4, 5, 7, 8, 9, 11, 12, and 13 (Sup. table 5), and identified the likely location of these SNPs relative to nearby genes using *SNPdat* (Doran and Creevey 2013). Additionally, we observed that the 2^nd^ most significant SNP in region 12, and the 2^nd^ and 3^rd^ most significant SNPs in region 11 were distinct from the surrounding SNPs, and resided in genes. Looking at these 13 SNPs, five SNPs were intronic, five SNPs were in the coding sequence, and three SNPs were intergenic.

We found no examples of the generation of a premature stop codon or a synonymous CDS substitution in these 13 SNPs, but several examples of coding sequence changes. Regions 4 and 5 each had a coding change, and regions 11 and 12 each had a coding change at the second-most-significant SNP in their peaks. Region 11 additionally had a nonsynonymous mutation at its third most significant SNP. These five SNPs, respectively, reside in genes that align in a *BLAST* search to the following genes: *rumpelstiltskin, pyridoxal kinase, CG10413, CG7627*, and *branchless*. Some of these gene orthologs have essential functions that could be associated with major phenotypic effects in mutants: *rumpelstiltskin* is associated with mRNA 3′-UTR binding and axis specification in embryos (Jain and Gavis 2008), and *pyridoxal kinase* is the enzyme that produces metabolically active vitamin B6 (Meisler, Nutter, and Thanassi 1982). Perhaps most interestingly, *CG10413* is associated with sodium-potassium-chloride transport, which is likely to be important to fitness in a vernal pool organism because of the variable salinity of a drying pool of water. We tried aligning the most promising genes, *CG10413* and *rumpelstiltskin*, to human, *D. melanogaster, Caenorhabditis elegans*, and *Mus musculus* orthologs identified via Flybase using the T-Coffee suite (Notredame et al. 2000). They showed no evidence of highly conserved protein domains, but because we were comparing them to very distant relatives, this does not rule out the possibility of conservation of their domains in comparisons with more closely related species.

In several cases the handful of SNPs driving the signal of a peak are associated with one or two populations harboring outlier allele frequencies relative to flanking regions. For example, at the most significant SNP for region 11 (Fig. 4, C7, R11 lower panel) the allele frequency of the LTER population is a clear outlier; it appears that LTER is driving the signal in this region. Indeed, all of the populations except LTER have an allele frequency of zero at this SNP. A manual examination of the alignments of the Illumina data does not reveal any evidence of poor alignment or repetitive elements in this region. On the other hand, the second-most-significant SNP appears to have a large amount of variation in the allele frequencies between the populations, with no evidence that the LTER population differs dramatically in allele frequency from the other populations. Because *X^T^X* takes into account the relationships of the populations, it is possible for it to identify differentiated regions that would be missed by directly examining the allele frequencies. Thus, we cannot conclude from allele frequencies alone that LTER is driving the association. It is of note that we never see long, dozen or more, SNP haplotypes in these figures. This is consistent with linkage disequilibrium extending over short distances in this species, and the strength of selection acting on these regions being relatively small relative to recombination.

### Associations of population differentiation and environmental variables

When the clam shrimp were collected, up to 24 environmental variables (Sup. table 6) were recorded for each population. These run the gamut from geographic data (latitude, longitude, elevation), through abiotic ecological variables (pond size, pH, etc.), to biotic ecological variables (the presence of other shrimp species, the percentage of males in the population). We collapsed several of the variables highly correlated across the 11 ponds of this study (Supplementary figure 11). We further generated a set of “dummy” environmental variables corresponding to each of the study populations. For example, the “LTER” dummy variable would have a value of 1 for the LTER population and a value of 0 for all others. We then correlated each dummy variable with each environmental variable and further collapsed environmental variables with an absolute *ϱ*greater than 95% with a population dummy variable. The logic here was to avoid attributing a region associated with an environmental variable to that variable, in cases where that variable is statistical confounded with a single pond. This data reduction algorithm resulted in the following collapses: *Thamnocephalus platyurus, Streptocephalus mackeni*, and Cladoceran presence/absence were found to be identical, and highly correlated with longitude and WAL (thus collapsed to WAL); *Eocyzicus* presence/absence was collapsed to Ares; Tadpole shrimp presence/absence collapsed to Forsling; volume and surface area collapsed to LTER. Finally, the surface area-to-volume ratio and depth were collapsed; that said, because they do not display any significance peaks indicating regions of local adaptation (see below), they have been excluded from our figures We did not include the absolute number of males or hermaphrodites in our analysis because they are a simple function of the amount of soil hydrated in the laboratory, although we did consider the frequency of males. This reduced the number of environmental variables from 24 to 13. The final set of environmental variables is listed in supplementary table 7.

We used *Bayenv2* to generate Bayes factors at each SNP, for 11 dummy variables and 13 collapsed environmental variables (see supplementary table 7 for a complete list). Bayes factors differ from *X^T^X* in that they are only elevated when the allele frequencies at a SNP are correlated with the environmental variable in question. For each ecological variable, the Bayes factors (S. N. Goodman 1999) compare two hypotheses: either that the observed allele frequencies are due to ancestry alone, or that they are due to a combination of ancestry and selection that is correlated with an environmental variable of interest. Our Bayes factors were elevated if the “selection” hypothesis was more likely than the “ancestry alone” hypothesis.

We began our analysis of the Bayes factors by subjecting them to the same Q-Q plotting test that we used with *X^T^X*. We present here Q-Q plots for a single Bayes factor (association with collection date, Figure 5, panel A), because other Bayes factors had nearly identical Q-Q plots. We find that the empirical values and the gamma distribution fit to the empirical values are fairly linear, with a very slight ‘hockey stick’ signal at the upper end of the distribution. This effect is made less obvious because of the double-log of the data, but is indicative of a very dramatic increase in the slope when the double-log is reverted (not shown). Our simulated neutral dataset allows us to establish that a log10(Bayes factor) single SNP threshold of 4.97 holds the genome wide false positive rate to be 0.36. This threshold obtained via simulation agrees with the observed inflection point in the Q-Q plot. Note that, to avoid logging negative values, or fitting a gamma distribution to a distribution with negative values, the double-logged Q-Q plot was transformed by the following equation: log10(log10(Bayes Factor) + 0.76) + 2.032, such that the log10(BF) value of 4.97 corresponds to a value of 2.79, approximately the point in the Q-Q plot (Fig. 5) where the hockey stick effect begins.

**Figure 5:**
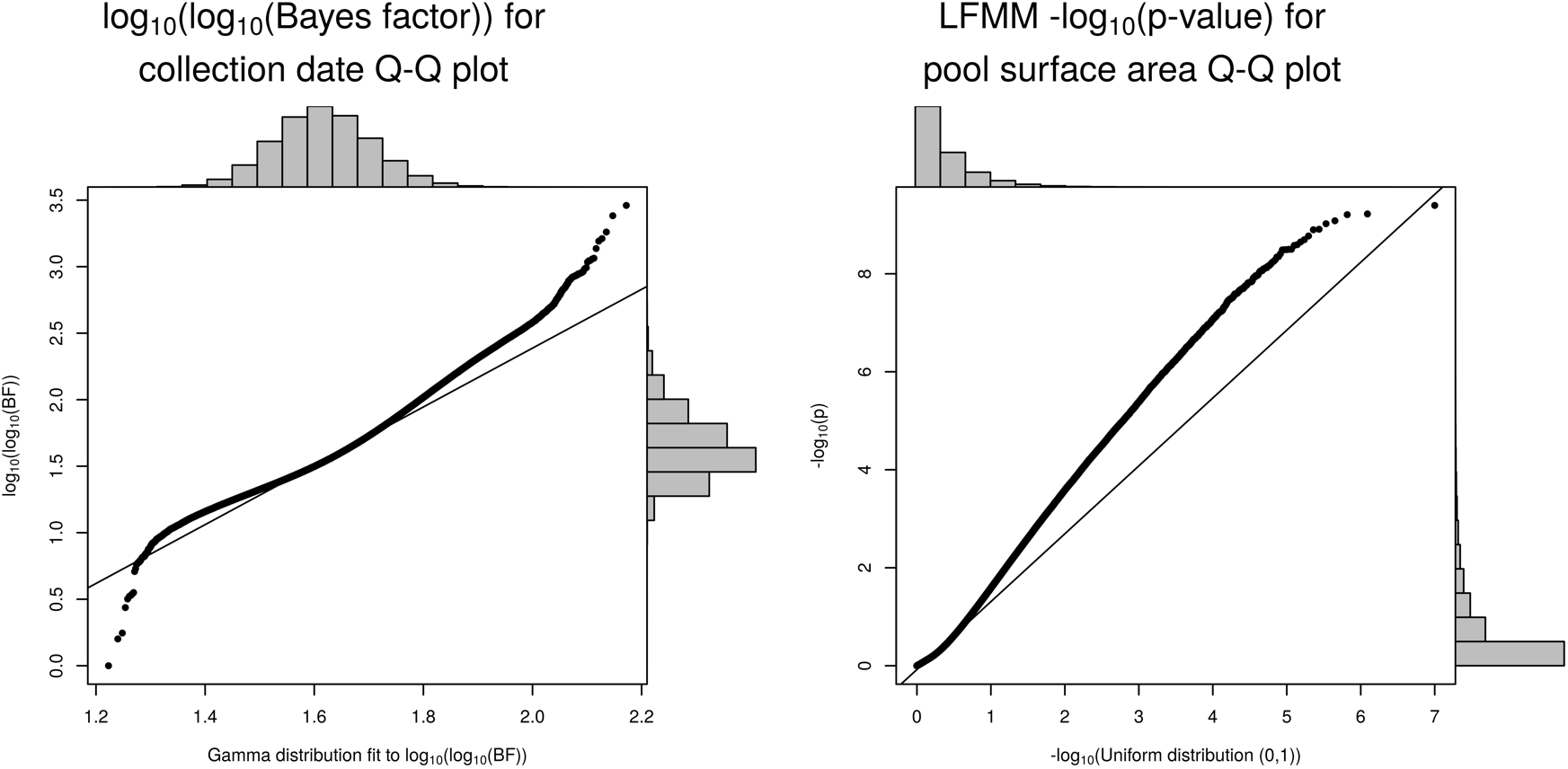
Quantile-quantile plots of two statistics used to identify regions with high population differentiation associated with collection date (an environmental covariate). Left: Bayenv2’s Bayes factors, double-logged, and plotted against a two parameter gamma distribution fit to the middle 70% of the distribution (methods) Right: LFMM’s negative logged *p*-values plotted against a negative logged uniform distribution from 0 to 1. The straight line is the linear regression line to the middle 70% of the data in both cases. The bar charts along the sides of the graph are histograms of the data along their respective axes.

We also discuss an alternative to Bayes factors, *LFMM*’s z-values (Frichot et al. 2013), but only in limited detail. *LFMM* does not take into account coverage, and so seems to produce inflated values in regions where coverage is low and allele frequency estimates are inaccurate (data not shown). Again, we begin by generating Q-Q plots of all *LFMM p*-values, though for the sake of brevity, we only report one sample plot here, as all others were similarly shaped. Our Q-Q plot of the -log(*p*-values) produced by *LFMM* for the environmental variable of collection date (Fig. 5, panel 2) against a negative logged uniform distribution between 1 and 0 (the expectation for a *p*-value), displays an upward “hockey stick” in the upper portion of the distribution (the majority of the data is in the very low end of the -log(p) value range, so a large portion of the Q-Q plot makes up the “hockey stick”). Genome wide analysis of our data with *LFMM* produced conclusions superficially similar to the Bayes factor results, which at first seems to be in keeping with the Q-Q plot results: a relatively small number of regions had visibly large numbers of strongly significant SNPs adjacent to each other. On the other hand, while a few *LFMM* hits seem to correspond to Bayes factor hits, many hits are unique to one of the two methods. Where there is disagreement, we believe *Bayenv2* is a more reliable indicator of the presence of local adaptation. *Bayenv2* incorporates count data into its significance calculations, while *LFMM* uses only allele frequencies. Comparison to coverage indicates that many of *LFMM’s* strongest hits are in areas of low sequencing coverage. This is unsurprising, as inaccurate estimates of allele frequencies should produce allele frequencies that are not in agreement with the existing relationships between the populations (Fig. 4). For this reason, we do not further discuss the LFMM peaks.

Across all 13 of our collapsed environmental variables (Sup. Table 7) we found 1,414 SNPs with at least one significant Bayes factor; similarly, across the 11 dummy variables (Sup. Table 7) we found 3,877 such SNPs. 1,520 genes are associated with at least one significant SNP via the environmental variables, while 3,504 genes are associated with at least one dummy variable. The Bayes factors thus seem to have tagged a larger fraction of the genome than the previous analysis that ignored environmental predictors and only employed *X^T^X*: *X^T^X* identified a total of 510 SNPs; Bayes Factors, all told, identified 5291 SNPs. We will discuss this difference in the next section. Unsurprisingly, when we censor the SNPs to only include those in the 13 regions, we find the same number of associated genes (50) as we do with the *X^T^X* analysis earlier.

The difference in the number of SNPs between the *X^T^X* and Bayes Factor analyses is due primarily to the Tank011 dummy variable, and the Elevation environmental variable. Both have extremely large numbers of significant SNPs (2,944 for Tank011, versus 92.3 for the average dummy variable). This difference in the number of SNPs could be due to either an incorrectly set significance threshold for these two tests, or it could be due to an especially high number of truly differentiated SNPs associated with these tests. The fact that Tank011 is one of our most deeply sequenced populations and happens to have a large number of significant loci seems to be circumstantial evidence that these may be the result of true signals of differentiation, but this is uncertain; Therefore, we focus here on the 13 large peaks that were supported both by *X^T^X* and a large number of Bayes Factor tests.

We next generated Manhattan plots of the Bayes factors for each of the 13 environmental variables (Figure 6, Figure 7, Sup. Figures 5-10) and 11 dummy variables (Sup. Figure 12). Because of the lack of peaks associated with some variables, figures depict only those variables that had at least one visible peak. Figure 6 provides an example of Bayes factors for the effect of collection date. As the shrimp populations were samples in different years (1995, 1996, 1998, 2000, and 2003), collection date presumably reflects differences in allele frequency due to the ecological details of the prior spring’s rainy season. Here we observe 3 peaks. Interestingly, these peaks correspond to numbered regions 3, 9, and 11 of Figure 7. An important observation is that we do not identify any additional peaks using Bayes factors and the environmental variable date that are not identified using the 11-population *X^T^X* test alone. A similar trend holds for the other 8 environmental covariates shown (Fig. 7). In many cases we identify peaks, but they always correspond to the peaks from the 11-population comparison. The same pattern holds for the 11 dummy variables (Sup. Fig. 12). Whenever the Bayes factor identifies a clear peak, that peak always corresponds to one of the 13 peaks identified using the *X^T^X* test. Further, because we cannot differentiate especially well between hypotheses 4 and 5 (above), there are reasons to believe single SNPs identified as significant that are not associated with a peak are simply false positives. Thus, we focus our analysis on the 13 peaks that seem most reliable.

**Figure 6:**
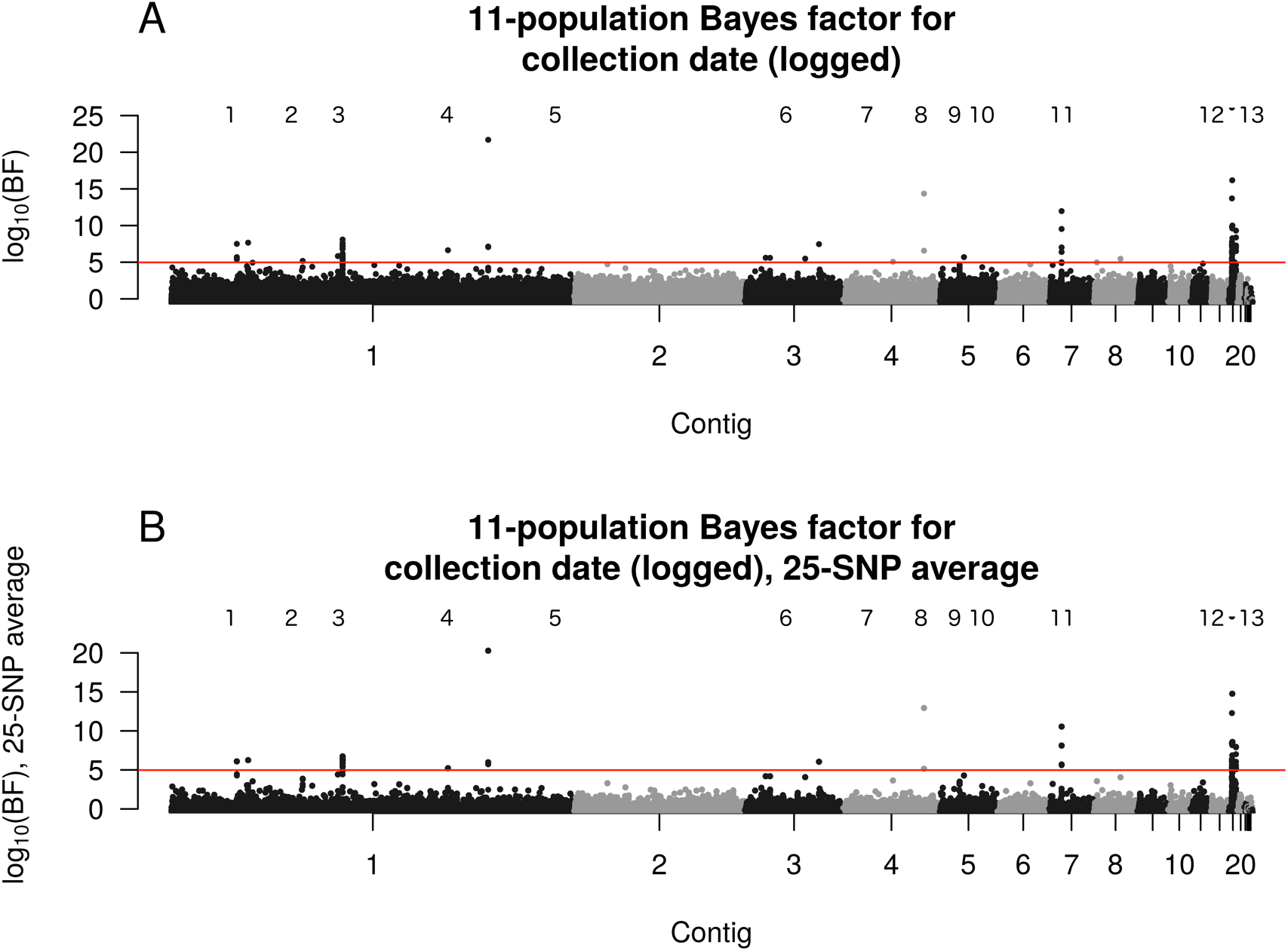
Manhattan plots of tests for an association between collection date and allele frequency difference for the 11 populations. A: Single SNP Bayenv2 Bayes Factor, logged; B: 25-SNP windowed Bayenv2 Bayes Factor, logged. Peaks numbered as in Figure 5. Red lines correspond to significance thresholds set as described in the methods.

**Figure 7:**
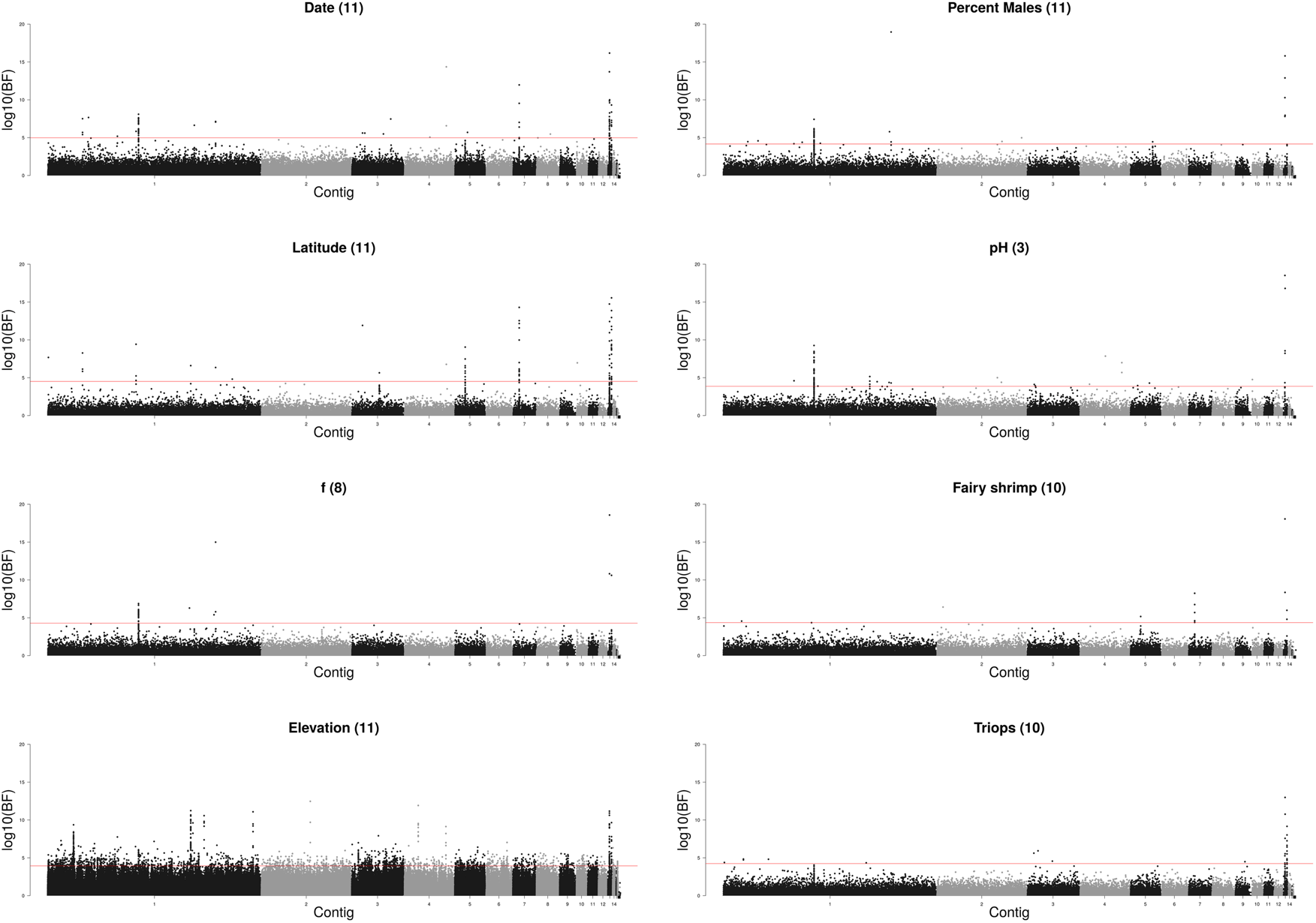
Manhattan plots of single SNP Bayes factors for different environmental variables. All plots show the 11-population logged Bayes factor associated with the given environmental variable, with environmental variables indicated, and the number in parenthesis the number of populations that environmental variable was measured for. “f” refers to the inbreeding coefficient estimated for each population (S C Weeks and Zucker 1999). Environmental variables that are highly correlated with a single pool, or that have no significant peaks besides regions 12 and 13, are not shown.

A summary of the mapping of peaks to environmental predictors or dummy variables is presented in Figure 8, with some environmental predictors and dummy variables excluded due to a lack of peaks associated with these variables. We searched for trends in the patterns of peaks relative to environmental and dummy variables. Most strikingly, regions 12 and 13 display elevated Bayes factors for every environmental variable we have measured. The putative genes identified in these regions (Cp1, CG7627, CG4562, multidrug resistance-like protein 1, octopamine β-1 receptor), which relate to various nervous system functions and wound healing, may be worth further inspection. That being said, peaks 12 and 13 are located in a small contig with somewhat high estimates of residual heterozygosity in the reference genome. A small assembly contig may be a red flag that the contig contains repetitive elements that would make alignment to the contig less reliable (Baldwin-Brown et al. 2017, Sup. Fig 3 – 14th contig from the left). Indeed, we see large regions of this contig that contain no SNPs that have passed coverage filtering (not shown), indicating that alignment to the contig in question is not as reliable as alignment to other contigs. Thus, we believe these peaks should be treated with some skepticism and may not be reliable. In this way, these regions highlight the value of a high quality genome assembly: alignment to a higher quality assembly is much more reliable.

**Figure 8:**
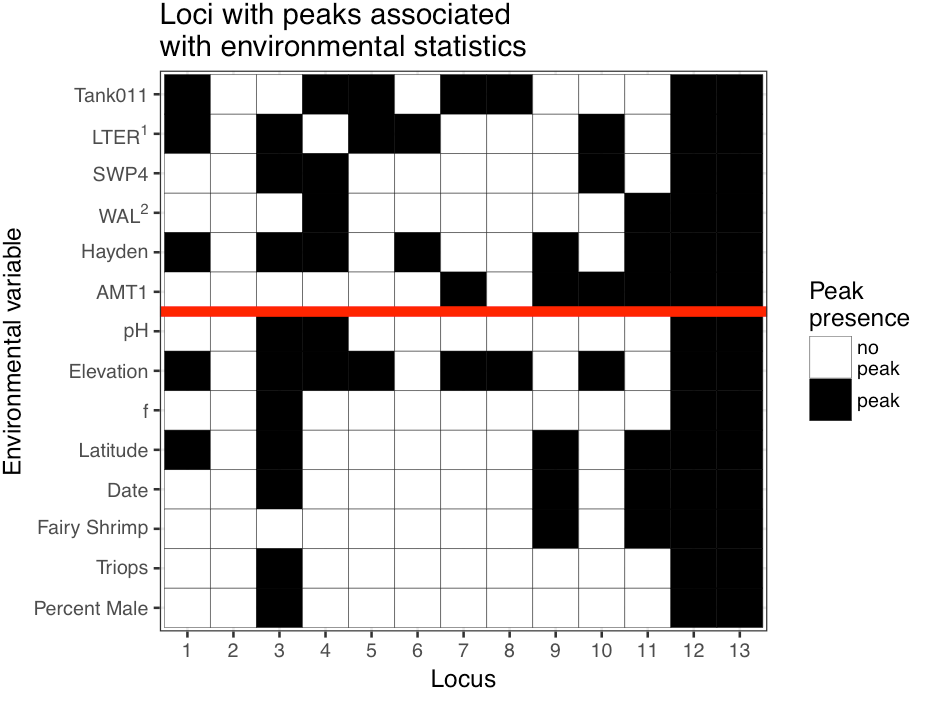
A matrix associating the regions of Figures 3, 6, and 7 with environmental variables or single populations that identified the same regions. A black square indicates the presence of a peak at that locus, when associations are tested using Bayes factors for that variable. A white square indicates no peak in that region, for that variable ^1^LTER is highly correlated with the surface area and volume of the pools. ^2^WAL is highly correlated with longitude, as well as the presence/absence of Cladocerans, S. mackeni, and T. platyurus. Variables above the red line are population dummy variables, while variables below the red line are measured environmental variables. Region 2 was not observed as correlating to any of the environmental variables or population dummy variables tested. All variables that were correlated with only regions 12 and 13 were excluded from this matrix.

The other 11 peaks are associated with regions of the genome with excellent assembly properties. Collection date is associated with regions 3, 9, and 11, and is of note because it is not highly correlated with any particular population. Of note here is the fact that collection date is not necessarily identical to the year of last hydration in the wild, as not all pools are filled with water on a consistent yearly basis. Collection date is not necessarily identical to the year of last hydration in the wild, as not all pools are filled with water on a consistent yearly basis, but is certainly connected to it. Elevation is associated with numerous regions: 1,3,4,5,7,9, and 10, and is also not correlated with any single population. It still remains a challenge to know if the observed associations with environmental predictors are real, or just happen to be associated with a subset of our 11 populations due to sampling.

With the above caveats in mind, some gene functions do seem to suggest a relationship between genotype and phenotype: for example, CG10413, a gene in region 11, is believed to have sodium/potassium/chloride symporter activity, and contains a nonsynonymous SNP with a very high *X^T^X* value; one might speculate this influences salinity tolerance in these shrimp as ponds are likely to vary in their salinity. Region 11 is associated with the collection date, the latitude, the presence of fairy shrimp, the AMT1 population, the Hayden population, and the WAL population (which is correlated with longitude, the presence of *Streptocephalus mackeni*, and the presence of *Thamnocephalus platyurus*). With so many associations, it is not likely that we can definitively identify the environmental variable that drives differentiation of CG10413. Likewise, *rumpelstiltskin*, a gene in region 4, is a development gene in *D. melanogaster* associated with embryo segmentation and anterior/posterior differentiation, and contains a nonsynonymous mutation at the SNP with the highest *X^T^X* value in region 4. We do not find evidence that this SNP is in a conserved residue, but our ability to identify conserved residues is limited by the lack of available genomes that are closely related to *E. texana*. Unfortunately, *rumpelstiltskin* is associated with elevation, pH, and the Hayden, WAL, SWP4, and Tank011 populations, so it is difficult to draw a conclusion about its relationship with any specific environmental variable. Other genes with nonsynonymous mutations in highly differentiated SNPs in the regions assayed here include *pyridoxal kinase* in region 5 (vitamin B6 production), and *branchless* in region 11 (branch morphogenesis in trachea and lungs).

## Discussion

### Wild populations and selection

Numerous studies have used an analogue of *F_ST_* for detecting local adaptation in model systems with a high quality reference genome with a large number of individuals genotyped. Studies performed in humans (Mackinnon et al. 2016), *Drosophila* (Reinhardt et al. 2014), and other model organisms (McGaughran et al. 2016) commonly use either whole genome sequence data (*Drosophila*, other models) or relatively carefully ascertained SNPs from genotyping chips (human). Given a high quality reference genome and large population sample, the identification of *F_ST_*-like outliers seems to be a somewhat effective means of detecting local adaptation (Savolainen et al. 2013). Only a small number of studies in non-model systems have employed a high-quality reference and genotyping dataset (e.g., Lamichhaney et al. 2015, McGaughran 2016). Perhaps the best non-model example at this time is Lamichhaney 2015 (Lamichhaney et al. 2015), a project that sequenced 120 finches across the Galapagos Islands using whole genome sequencing, and used a high quality (5.2Mb scaffold N50) assembly for alignment. In spite of the fact that they restricted themselves to a normalized version of *F_ST_*, which does not account for relationships between populations, they were still able to detect several regions that they believed contained genes related to beak morphology, a trait that did not escape Darwin’s attention. Even so, many of these peaks were about the same size as the scaffolds on which they resided, making it difficult to tell where one peak ends and another begins. McGaughran 2016 (McGaughran et al. 2016) sequenced 264 strains of nematodes *(Pristionchus pacificus* rather than *Caenorhabditis elegans)* and performed an *F_ST_* analysis, the authors successfully identified a set of locally adapted regions. One region contained a gene that was a homologue of the *C. elegans NHX* gene family. They confirmed this gene’s effect on pH tolerance using transgenics. Both of these studies required sequencing of a large number of individuals from a broad geographic area, and the construction of a high qualify reference genome. Both were successful at identifying regions important in local adaptation using *F_ST_*.

In contrast, scans for local adaptation in non-model systems are typically performed under less than ideal conditions. We could not identify any examples in the literature of a high quality genotyping dataset associated with highly fragmented reference, nor studies with low quality genotyping but a very good reference. Typically, non-models have highly fragmented references and relatively sparse genotyping datasets. A low contiguity reference genome assembly can prevent researchers from being able to identify peaks of significance – if a peak is larger than the contig that contains it, it is possible that the multiple sections of the peak will be identified as separate peaks because they are on separate contigs. Even in the case of a relatively high quality assembly (i.e., Lamichhaney et al. 2015), this problem can still occur in the smaller contigs. The worst possible instance of this would be a complete lack of a reference, as in Lal et al. 2016, where SNPs can only be analyzed independently, and there is no concept of a significance peak. Furthermore, in our work two of our thirteen peaks are likely artifacts as they are associated with contigs of dubious quality; it is unclear the extent to which a high fragmented assembly would magnify this source of false positives. On the other hand, a small number of assayed markers can lead to the problem of missing all of the SNPs in a region of population differentiation completely or sampling SNPs too sparely to detect a peak. In our study many peaks were only tagged by a few dozen SNPs; we might expect this to be the case in other systems in which linkage disequilibrium only extends over short physical distances. There are many examples of non-model systems that have had populations sequenced using candidate gene sequencing (Keller et al. 2012), RADseq (Lal et al. 2016), targeted genomic sequencing (Yeaman et al. 2016; Roulin et al. 2016; Holliday et al. 2016), and other methods (Riginos et al. 2016; Wenzel et al. 2016). Although these techniques for ascertaining polymorphism information are known to be reliable and affordable, most of them are limited in the number of polymorphisms that they can assay (often as low as a few hundred loci). For comparison, we used a total of 1.4 million SNPs for our *X^T^X* scan, giving us an average resolution over the 120Mb genome of one marker every 85 basepairs. A set of only 500 SNPs on this same genome would give a resolution of one marker every 240kb, leaving large gaps of unsampled genomic content that could prevent the detection of population differentiation if such differentiation is localized to a small region.

In most published studies, *F_ST_*-like statistics are computed using population-wide allele frequency data. Most use *F_ST_* or a derivative thereof (Lamichhaney et al. 2015; McGaughran et al. 2016), but others use more complex statistics such as those of *Bayenv*, (Keller et al. 2012), *Bayescan* (Foll and Gaggiotti 2008; Keller et al. 2012; Lal et al. 2016), *LOSITAN* (Antao et al. 2008; Lal et al. 2016), or others. We used *F_ST_*, *Bayenv2*’s *X^T^X* and Bayes factors, and *LFMM’s* z-values to identify signals of selection in these populations. We largely disregarded *F_ST_* in our study because there are clear relationships between the populations that make *F_ST_* poorly suited to identify local selection. Indeed, we found that we could not distinguish any peaks of localized population differentiation using *F_ST_*. Additionally, we largely disregarded the results of *LFMM* because it fails to take into account coverage when computing significance, and many of *LFMM*’s peaks appear to occur in areas of suspiciously low coverage, where estimates of allele frequencies are inaccurate. It is reasonable to assume that *LFMM* should be very effective in scenarios where the allele frequencies are known to have been accurately measured (say, a large sample of individuals that are individually sequenced rather than sequenced in pools and/or genotyped using SNPchips), but when the sequencing coverage used to estimated allele frequencies varies *LFMM* appears unreliable. Therefore, we relied largely on the results of *Bayenv2*’s *X^T^X* and Bayes factor statistics when dissecting this data. *Bayenv2* identified a number of regions of localized population adaptation, but is extremely computationally expensive. The Markov chain Monte Carlo approach to parameter estimation requires at least thousands of iterations per locus, and each locus must be estimated multiple times (five, in our case). On top of that, identification of a significance threshold requires further *X^T^X* and Bayes factor estimation on simulated neutral datasets, which must be considerably larger than the genome. At ~5 minutes per run per locus on a modern high performance cluster, we estimate this project required greater than one million CPU hours. A similar project, performed on a human-sized genome, or one on the same sized genome with an order of magnitude more populations would be computationally difficult to carry out. The most likely reason the Bayenv2 approach has not been used on genome wide datasets consisting of millions of SNP markers is simply that of computational difficulty. The field would be greatly accelerated if future versions of *Bayenv2* were ~100X more efficient.

To our knowledge ours is the first non-model study employing a high contiguous genome assembly, a high coverage genome wide Poolseq dataset that detected millions of SNPs in several populations, and a computationally challenging, but powerful, Bayesian statistic framework. The intersection of assembly, whole genome sequencing, and complex Bayesian statistics in our case led to the detection of regions of the genome apparently under local adaptation that would not have been detectable using simpler methods, perhaps owing to the relatedness of some of our surveyed populations and the relatively small number of populations surveyed. It is difficult to directly compare our study to others, as nearly all of these studies relied upon sequencing of individuals, rather than sequencing of pooled populations (with rare exceptions J. Chen et al. 2016). Our use of pooled population sequencing allowed us to estimate our allele frequencies based on a large total number of individuals, but also caused uncertainty in the number of individuals contributing to an allele frequency estimate, and caused the number of individuals contributing to each estimate to differ between both from region-to-region and from population-to-population. Thus, it is difficult to compare the value of the 844X of Illumina sequencing performed here across all 11 populations to, say, the 120 individuals sequenced in (Lamichhaney et al. 2015).

### Wild populations and selection in *E. texana*

Conventional wisdom indicates that vernal pool organisms are not capable of a great deal of migration under most circumstances. Indeed, Bohonak 1998 indicates that a geographic distance of only a few hundred meters should be sufficient for a high degree of differentiation of populations in the Anostraca (fairy shrimp). The inability of vernal pool shrimp to escape the pools in which they are born seems to prohibit migration between distinct pools. In fact, we find here (Fig. 1) that there is a great deal of migration between pools, both across short and long geographic distances, with shorter distances leading to increased migration - mean pairwise *F_ST_* is 0.038 across these samples. The source of this ability to migrate, whether it is animal tracking, wind dispersal, periodic flooding, or some other mechanism, should be the subject of further study.

We identify several genomic regions that appear to be under selection, as well as several variables in the environment that appear to be correlated with these selected regions. Of note are two regions that appear to be subject to selection that are both related to RNA-to-protein translation, including *CG10306*, which is expected to be involved in regulation of translation initiation (FlyBase Curators, Swiss-Prot Project Members, and InterPro Project Members 2004), *La*, which is experimentally validated as binding to rRNA primary transcript (Yoo and Wolin 1994), and *rumpelstiltskin*, which is experimentally validated as binding to the 3’ UTR of mRNAs. It is not clear what would drive protein translation machinery to be under differential selection in different pools, though we note that not all genes in our assembly have orthologs in *D. melanogaster*, so it is possible that unannotated genes, or even undiscovered genes, could drive these signals of local adaptation.

Correlation with environmental variables seems to indicate that certain variables have a larger effect on allele frequencies than others, but there are good reasons to think that the associations with environmental variables that we find are not proof of a connection between the environmental variable and adaptation. For example, there are a number of regions (1,3,5,6,10,12, and 13) whose allele frequencies are strongly correlated with surface area of the pool in which the shrimp reside. Pool surface area could easily be a strong agent of local adaptation, including the persistence of water over longer periods, the presence of predatory shrimp such as tadpole shrimp (there may be a relationship between pool size and presence of tadpole shrimp), consistency of food availability, etc. Unfortunately, only a single population (LTER) is represented by a large pool, so peaks associated with pool area are confounded with the LTER population, and it is difficult to draw conclusions about pool area without a more extensive sampling that includes both large and small pools. A better example of an interesting environmental predictor is collection date. All samples in this data set were collected in one of five collection trips in the years 1995, 1996, 1998, 2000, and 2003. It is easy to imagine that selection pressures differ between, say, a rainy year and a dry year, and that changes in climate could drive temporary adaptation. Further work could determine if there is a relationship between the previous season’s rainfall and the allele frequencies at collection date-associated regions. Overall, although a large number of SNPs were significantly differentiated, a small number of regions showed visibly strong signals of local adaptation. These few, strongly differentiated regions certainly seem to be worthy of further study. The clam shrimp ortholog of CG10413 in region 11, for example, is predicted to have sodium/potassium/chloride symporter activity. It has long been believed (Potts and Durning 1980) that sodium/potassium pumps and chloride pumps with passive sodium diffusion are important for regulating osmotic stress in vernal pool shrimp. Although we have no data on salinity in these pools (and it likely various over the life of any given vernal pool), it is tempting to speculate that salinity is correlated with the significant region 9 hits. We identified five genes, *rumpelstiltskin*, *pyridoxal kinase, CG10413, branchless*, and *CG7627*, that have nonsynonymous coding changes produced by one of the top two SNPs in one of our 13 peaks. These genes seem especially worthy of further investigation.

Studies have historically had much higher power when comparing allele frequencies to environmental variables, rather than merely to one another, with some selected regions being identifiable only when examined in the context of a correlation with the environment. For example, Berry and Kreitman 1993 observed excess differentiation at two functional sites in the *Adh* gene in Drosophila relative to the latitude of the collection site, but were unable to detect the same two sites by just looking for excess differentiation between populations. In contrast, we found that we were able to identify all of the major environmental variable-associated peaks using only an ecology-agnostic measure of population differentiation (*X^T^X*). This seems to be a demonstration that modern statistical techniques, combined with whole-genome SNP discovery and analysis, have a much higher power to detect differentially selected sites without knowledge of the ecology of the organisms in question.

### The future

We identified a relatively small number of candidate genes that appear to be associated with differentiation of these populations. Genetic studies, perhaps using CRISPR-Cas (Jinek et al. 2012) could reveal much about the effect of these genes on phenotype. That being said, the approaches of this work only identify regions important in local adaptation - they do not tell us what phenotypes variation in these regions impact. So, even with the ability to generate gene knockouts, it is unclear what phenotypes should be examined. The observation that somewhat reproductively isolated populations living in different ecological or physical settings can show strong geographical isolation in isolated genomic regions suggests a powerful paradigm for identifying the genes contributing to adaptation in the wild. Further reductions in the cost of collecting Poolseq datasets would allow studies such as this to be carried out with many more populations, allowing the effects of specific populations versus environmental characteristics of those populations to be statistically disentangled. It would currently be a major computational challenge to efficiently calculate statistics such as *X^T^X* genome-wide for larger collections of populations. If that challenge could be met, future studies are more likely to be limited by the ability to measure a large number of relevant ecological properties for each population, rather than the ability to accurately estimate the allele frequency at every SNP genome-wide. The methodologies of this paper provide a blueprint for characterizing the genetics of local adaptation in never-before-sequenced species.

## Acknowledgements

This work was supported by NIH grants AI126037, GM115562, and OD10974 to Anthony D. Long, and NSF grant DEB-9628865 to Stephen C. Weeks. Thanks to Stuart MacDonald for sequencing at the University of Kansas.

## Data Availability

All sequencing data is available at the NCBI All data will be made available at the NCBI Sequencing Read Archive under the Bioproject “PRJNA352082”. Additional files are available at the following URL: http://www.wfitch.bio.uci.edu/~tdlong/PapersR_awData/BaldwinShrimp.tar.gz. Additionally, all scripts used for analysis will be made available at the following GitHub page: https://github.com/jgbaldwinbrown/jgbutils

## Author Contributions

Experimental design by JB and AL. All analysis and manuscript drafting performed by JB. Revisions by JB and AL. Shrimp rearing, DNA extraction, and sequencing library preparation by JB.

